# Local resource heterogeneity drives density-dependent dispersal expression in a phoretic mite

**DOI:** 10.1101/2025.07.29.667385

**Authors:** Lukas H.A. Edwards, Isabel M. Smallegange

**Author notes:** Corresponding author: Lukas Edwards. **Author contributions** LE conceived the study idea, as well as planned and conducted the experiments, with study design being advised by IMS. LE analysed the data and wrote the manuscript, with advice and input from IMS. Both authors contributed towards the revision of the manuscript.

## Abstract

1. Dispersal is favoured in highly heterogenous and ephemeral environments where it can increase fitness. However, whilst we generally assume that unfavourable local conditions cue dispersal to elsewhere, little is known about how heterogeneity in local conditions influences dispersal expression. This is an important element of dispersal, since dispersal characteristics rely on such local cues to signal their development.
2. We assessed the role of local environmental heterogeneity on the expression of a plastic dispersal phenotype in experimental populations of the bulb mite *Rhizoglyphus robini* through a multigenerational, 18-week experiment. Populations were kept under either heterogenous, homogenous, or intermediate food distribution treatments, and at low or high population density, via culling. This allowed us to test for the effect of local environmental heterogeneity and competition on dispersal expression, but also on population size and growth.
3. We found that heterogenous environments induce greater expression of dispersal in high density environments, but not in low-density ones. Additionally, whilst culled treatments have on average larger population sizes than non-culled treatments, non-culled and heterogenous environments have similar sizes to culled treatments. Treatments did not impact population growth rate.
4. Our findings that heterogenous treatments only induce greater dispersal expression under non-culled treatments suggests that heterogeneity may exacerbate competition around feeding patches, causing infrequent access to food. We also infer that culled populations become larger than non-culled, at least in the short term, due to rapid compensatory growth causing an overrepresentation of juveniles in the population. It is unclear why unculled heterogenous environments exhibit a similarly large population size to culled environment but could be due to food being hard to access but easy to locate.
5. Overall, we find an ecologically significant impact of local heterogeneity on population size and dispersal expression, much of which may arise due to the influences of heterogeneity on competition around foraging spaces. Such findings highlight the need to distinguish between local and regional heterogeneity when modelling the evolution of dispersal, since these two alternate levels may perform significantly different roles in influencing the expression or adaptive value of dispersal.

## 1. Introduction

Dispersal is a key trait that can mitigate against inbreeding, reduce competition between siblings, facilitate gene-flow, and allow the colonisation of novel habitats (Hanski & Mononen, 2011). Additionally, at the individual level, dispersal can allow individuals to escape pests or competition for resources (Mehrparvar et al., 2013). However, dispersal is inherently risky and individuals may incur significant costs through all stages of dispersal, i.e. the pre-departure, departure, transfer, or post-settlement phases (Bonte et al., 2012). These costs can impact an individual’s metabolism, reproductive output and even mortality. Given these associated costs, dispersal is predicted to occur predominantly under suboptimal environmental conditions (Chapman et al., 2011; Bonte et al., 2012) where the potential benefits of locating a more suitable habitat are likely to outweigh the fitness costs of remaining in a deteriorating environment. Heterogenous resource distributions can cause intense competition at interspersed, local feeding patches (Vahl et al., 2007; Bowler & Benton, 2009; Bonte et al., 2012) and are thus an important driver of dispersal. But whilst heterogeneity at the larger, regional, between-habitat level impacts the transfer-stage of dispersal, we know little about the role of fine-scale, local heterogeneity on dispersal during the pre-departure and departure phases. Local heterogeneity is likely to play a critical role, as dispersal decisions are typically informed by local environmental cues, such as feeding rates and resource accessibility, which are themselves strongly influenced by the spatial distribution of resources (Lombaert et al., 2006; Bowler & Benton, 2009; Bonte et al., 2012; Cote et al., 2017; Seeman & Evans Walter, 2023). This raises questions such as how does local heterogeneity influence dispersal expression? What can we infer about the role of resource distribution in competition? And what does this tell us about the role of local heterogeneity in the evolution of dispersal?

This study aims to quantify the role of local heterogeneity in food conditions and population density, used here as a proxy for intraspecific competition, on the facultative expression of dispersal phenotypes prior to departure. To meet this aim, we conducted a long-term, multigenerational population experiment using the acarid bulb mite *Rhizoglyphus robini*, which has a facultative dispersive life-stage called the ‘deutonymph’ (Diaz et al., 2000; Seeman & Evans Walter, 2025). The deutonymph life-stage is expressed just prior to the final juvenile life-stage; it does not feed and develops morphology suited to phoresy (Díaz *et al*. 2000). Phoresy is a common form of dispersal among mites and other soil living invertebrates (Borges, 2022), in which individuals attach themselves to larger organisms like arthropods, to be carried to novel habitats before detaching and meta-morphing into the later life-stages (Borges, 2022; Seeman & Evans Walter, 2025). This dispersal strategy requires specialised morphological adaptions, such as a sucker plate on their dorsal side, a distinct red colouration, and loss of feeding ability (Diaz et al., 2000; Borges, 2022; Seeman & Evans Walter, 2023). The costs of expressing these morphological adaptations include delayed development time and reduced reproductive output (Bonte et al., 2012; Deere & Smallegange, 2023).

In a multigenerational experiment, we subjected 30 populations of *R. robini* to three alternate food distribution treatments and two density treatments over 18 weeks. Food distributions were either one clump of food (‘heterogenous’), four small food patches (‘intermediate’), or powdered food scattered across the population (‘homogenous’). Meanwhile low and high-population density was maintained through either regular culling of half the population or not-culling populations, respectively. Importantly, all experimental populations were given the same total volume of food; thus, only the distribution of food and abundance of individuals varied between populations (Figure 2). We recorded the total number of individuals of all life-stages and sexes, which we then used to calculate total population size, population growth rate between observations (3-day intervals) and relative deutonymph expression. We predict that deutonymph expression (quantified as the proportion of deutonymphs against other individuals who have similarly experienced the sensitive window for this life-stage: tritonymphs and adults) will be highest in non-culled populations with heterogeneously distributed food, because this creates the most limited access to food. Furthermore, we predict that low-density populations with homogenous food distributions will have the highest population growth rate, and high-density populations with heterogenous food will have the lowest population growth. We also predict that, due to the high degree of compensatory growth in fast life-history species (Tanner et al., 2019), culled populations will have the highest population growth thanks to high numbers of eggs and larvae.

## 2. Methods

### The bulb mite

*Rhizoglyphus robini* has a short lifecycle and can mature after 9 to 40 days (Smallegange, 2011) and has a longevity between 31 to 130 days (Diaz et al., 2000). The lifecycle of *R. robini* progresses across five to six life stages: egg, larvae, protonymph, facultative deutonymph, tritonymph and adult. All life-stages except the egg and larval stage are preceded by a quiescent phase (Figure 1; Deere & Smallegange, 2023). Interestingly, *R. robini* has a second polyphenism, in which males may either develop into a large weaponised fighter or small weaponless scrambler. However, this reproductive polyphenism is only visible at the adult life-stage, which is also when one can identify the sex of individuals by their morphology.

**Figure 1.**
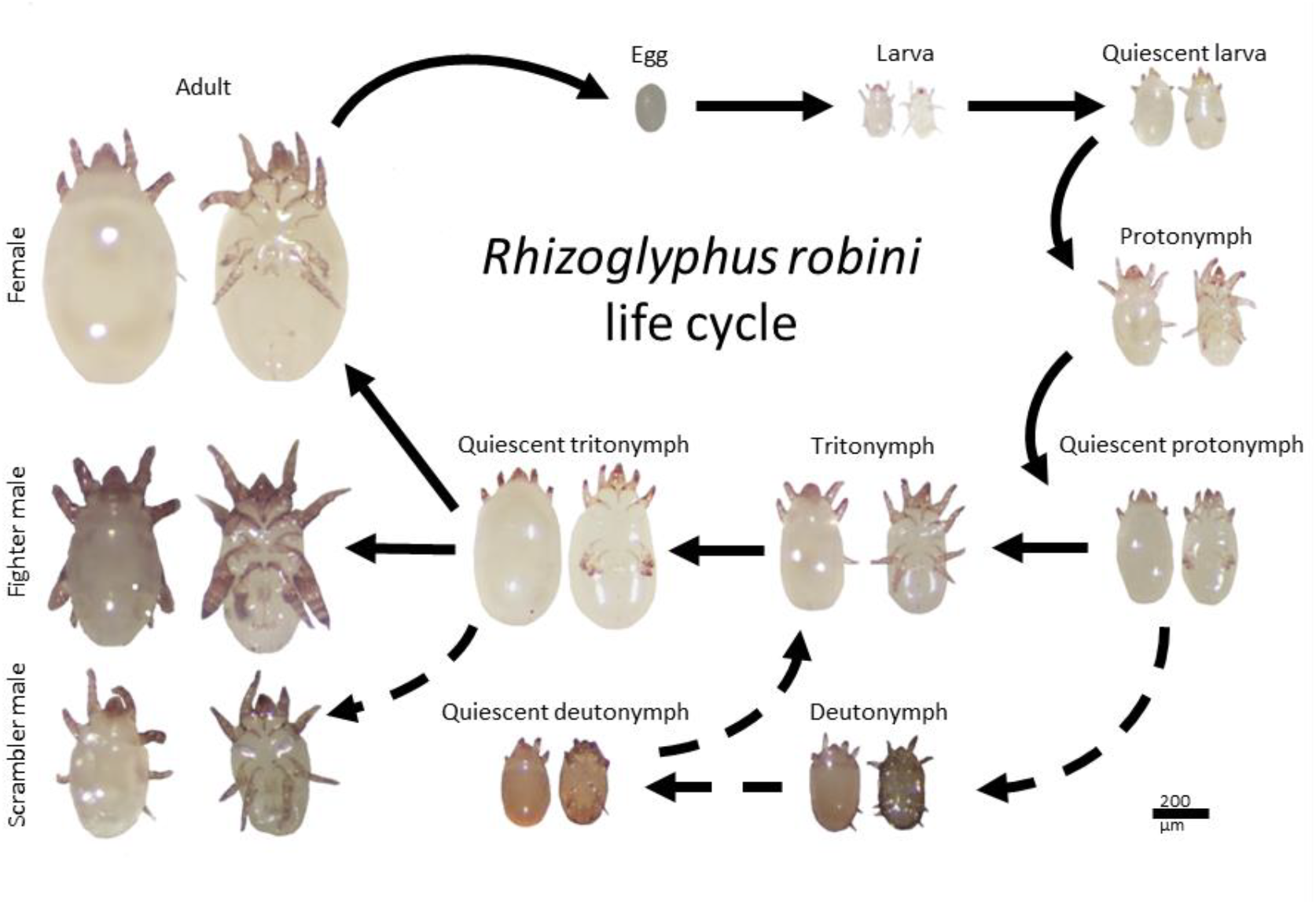
Life cycle of *Rhizoglyphus robini*. Obligatory developmental paths are shown as solid arrows, whilst plastic pathways are dashed. The life-cycle of *R. robini* has six distinct life-stages, five of which are obligatory and one which is expressed plasticly: the deutonymph. Notably, male morph is also a plastic trait hence the alternate fighter and scrambler males. This figure is largely taken from Deere and Smallegange 2023, Figure 2.

### Stock cultures and experimental populations

*Rhizoglyphus robini* were collected from flower bulbs in a garden in Newcastle-upon-Tyne in June 2023. Stock cultures were subsequently established in the laboratory with *R. robini* stock cultures maintained in incubators set at 25°C. Initially, the first 3 of these stock cultures had *ad libitum* to dried active yeast until February 2024 when another 4 cultures were started using individuals from the initial cultures. The new cultures were fed organic oats, a poor-quality food source often used to induce greater deutonymph expression (Deere et al., 2015), twice a week to ensure *ad libitum* access to food. Stock cultures were watered weekly, and cultures of a food type (i.e. cultures fed on oats or cultures fed on yeast) are mixed regularly to facilitate geneflow. Oat-fed stock cultures were allowed to acclimate to the new oat diet for three months prior to experiments, which equates to approximately 6 generations (Deere et al., 2015).

After acclimation, experimental populations were initiated from 100-200 individuals taken at random from the oat-fed stock cultures and placed into 30 experimental, 25 mm round glass population tubes. The lower third of the population tubes was filled with plaster of Paris, which was smoothed of any air holes to prevent burrows forming and the lids of the tubes had holes drilled in which were then covered in fine mesh, to facilitate air flow but prevent escape. After transfer, experimental populations were allowed to acclimate for a further 7 weeks, approximately 3 generations (2^nd^ May – 20^th^ June 2024), prior to experimentation.

### Experimental design

We started with 30 experimental populations, all of which were fed a food quantity equal to one large oat, twice a week. Ten populations were given one single, intact oat at the centre of the population (‘heterogenous’ treatment) (Figure 2). Ten populations were given a single oat but in four, equally sized quarters, one in each corner of the culture (‘intermediate’ treatment) (Figure 2). The final ten populations received a single oat that was crumbled finely and scattered evenly across the population (‘homogenous’ treatment) (Figure 2). Half of all cultures were randomly assigned to the low-density treatment and were culled by 50% once every two weeks (Figure 2), after counting, whilst high-density treatments were not culled. Treatment combinations were replicated 5 times.

**Figure 2.**
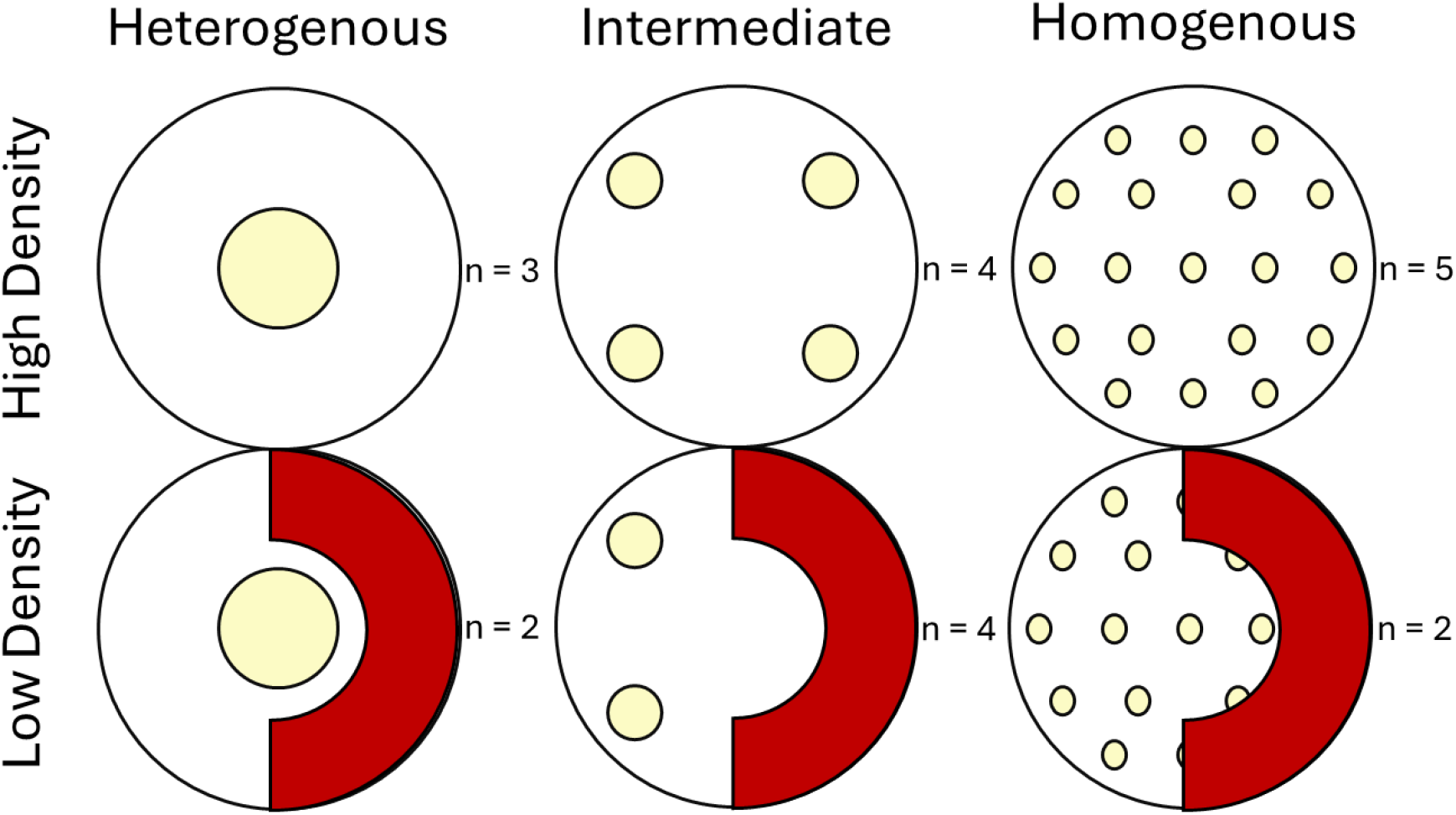
Experimental Design. A third of the experimental populations were given heterogenous (one whole oat), intermediate (4 quarters of an oat), or homogenous (one oat completely crumbled/powdered) food distribution, twice a week. Half of all populations were maintained at high density, and half at low density, by culling half the population every two weeks. For each treatment, *n* represents the number of populations which lasted the entire duration of the study.

We followed populations for 18 weeks, approximately 9 overlapping generations (Smallegange, 2011). Populations were observed visually through a Zeiss Stemi 2000-C microscope twice a week and the number of individuals at all life-stages and adult sex were scored with minimal invasiveness of one quarter of each population and subsequently multiplied by four to approximate total demographics (Tom Cameron, personal communication). After counting, we conducted the culling for populations in a culled treatment, if such culling was due. Finally, we fed and watered each population. Due to logistical problems, there was a two-and-a-half-week period (week 12-14 of the experiment) where observations were not made. Additionally, we had 10 compromised populations that did not perform well or died during the experiment because they had too much or too little water, leading to mould growth or dehydration, respectively. To account for any impact on populations that experienced such excess or lack of water, but which did not experience a population collapse, we recorded population status as either 0, meaning unperturbed, or 1, meaning perturbed. Our final dataset comprised 688 observations on 20 intact populations and 10 compromised populations (Figure 2).

## Statistical Analysis

We tested for any effects of food distribution (**F**; heterogenous, intermediate, homogenous), culling (**C**; non-culled, culled) and time period (**T**; binned 3-week intervals 1, 2, 3, 4, 5, 6) and all their two-and three-way interactions on deutonymph expression using a Generalized Linear Mixed Model (GLMM) with population tube and population status (intact or compromised) as a random factor. We used population status as a random effect since this is not expected to have a constant effect on the abundance of deutonymphs and is not a focal element of the study design. For this GLMM we used a beta-binomial distribution to account for overdispersion in the data. To model the binomial response variable, we used the cbind function to combine the counts of deutonymphs (successes) and number of non-deutonymphs (failures). To assess the effect of experimental treatments on population size we use a negative binomial GLMM, again to account for overdispersion. In this GLMM, we again used food distribution, culling treatment, time period (binned 3-week intervals), and all their interactions as fixed effects and population tube and population status as random effects on total population size. Finally, we use a linear mixed-effect model (LMM) to test the effects of food distribution, culling treatment, time period (binned 3-week intervals), and all their interactions on population growth rate. Population growth was calculated by dividing population size at an observation by the population size during the previous observation. This was done using the ‘lag’ function from the **tidyverse** library and controlling with population id. We also included population tube as a random effect within the LMM. Additionally, we performed a natural log transformation on the population growth rates to conform to the assumption of Gaussian error distributions.

We used model simplification to acquire the best model from the full model. The least significant term, starting with the highest order interaction, was first removed from the fitted model to produce a reduced model. If this removal led to a significant increase in deviance, we did not remove this term from the model and instead removed the second least significant term, again starting with the then highest order interaction. We tested for significant changes in deviance by using a likelihood ratio test. If a removal did not lead to a significant increase in deviance, then this term was removed from the model, and the next least significant term of the highest order interaction was removed. We repeated these steps until only significant terms remained in the model. We applied the same procedure to merge levels within categorical treatments that were not significantly different. Deviance significance was again tested using a likelihood ratio test. All analyses and model reduction was carried out using R (v4.3.0) (R Core Team, 2022).

## 3. Results

### Deutonymph expression

Deutonymph expression was significantly affected by the three-way interaction between food distribution, culling, and time period (T) (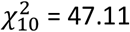, p <0.001). Within this interaction, deutonymph expression in the ‘intermediate’ and ‘homogenous’ food distributions did not significantly differ 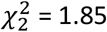, p = 0.3975) and were combined into one level, which we called ‘non-heterogenous’ food distribution. The model accounted for 91.7% of the variance, of which 73.6% is explained by fixed effects and 18.1% by random effects. The overall result was that the relative abundance of deutonymphs is lowest under non-heterogenous resource distributions or low population density but is highest when resources are heterogeneously distributed and population density is high (Figure 3). However, these interactions are much more pronounced during later timesteps, from T4 onwards, and there are no significant impacts of experimental treatment during timestep 1 or 2.

**Figure 3.**
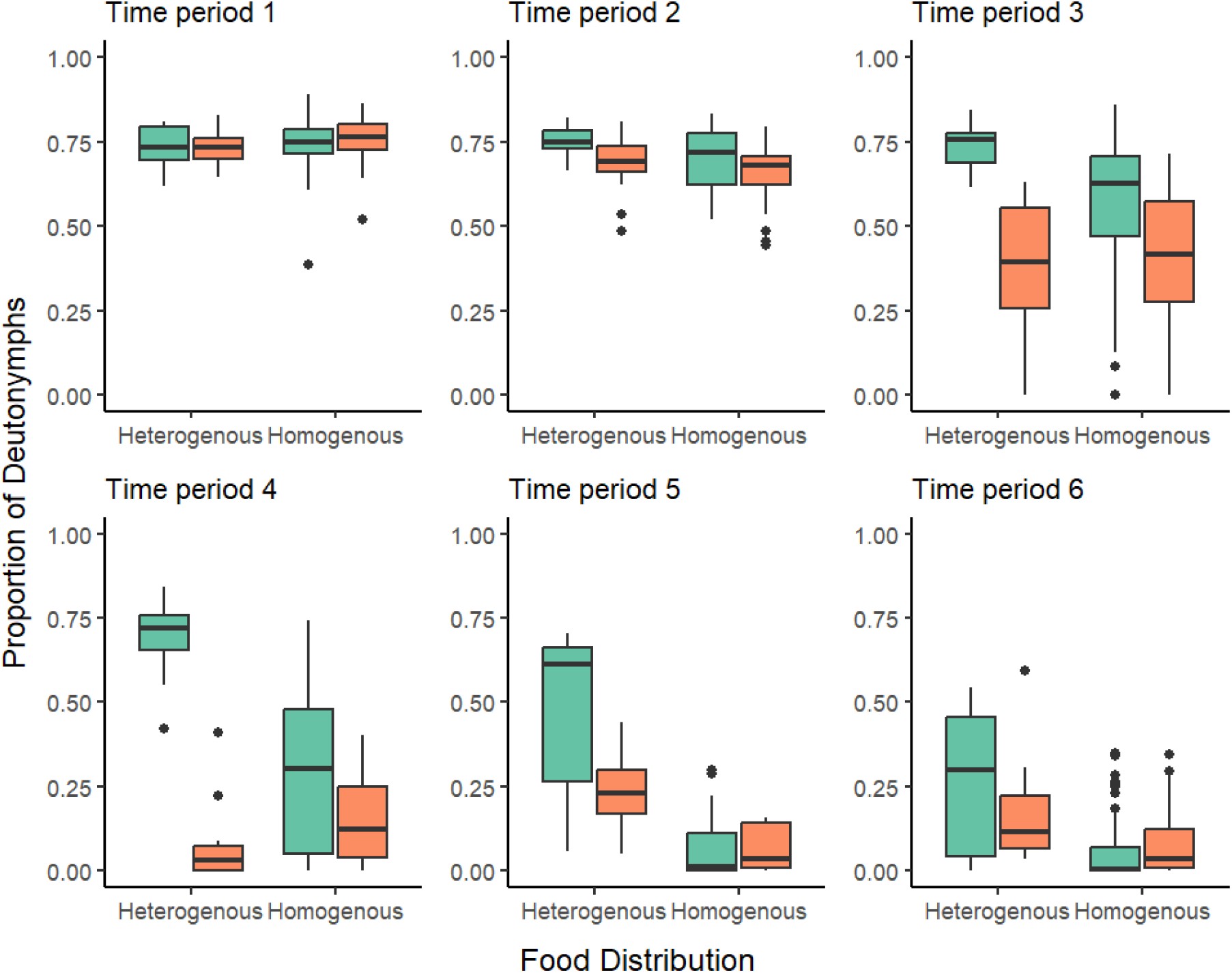
The effect of culling and environmental heterogeneity on the proportion of deutonymphs across time. Across time-blocks the proportion of deutonymphs within populations decreases significantly, with the earliest responses being in culled populations (orange) and slower responses from non-culled populations (green). Culled and non-heterogenous environments cue reduced plastic expression of deutonymphs, compared to the non-culled heterogenous environments.

### Population size

Population size is significantly influenced by the three-way interaction of food distribution, culling, and time period (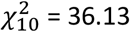, p < 0.001). Within time-period we found that T1 and T2 (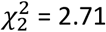, p = 0.844), T3 and T4 (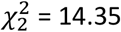, p = 0.279), and T5 and T6 (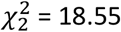, p = 0.420) could be merged without significantly increasing the deviance of the model. This creates three time periods, ‘early’, ‘middle’, and ‘late’, which corresponds to weeks 1-6, 7-20, and 13-18, respectively. This GLMM explains 74.5% of the variance. However, the majority of this (52.9%) is explained by the random effects, and only 21.6% is explained by fixed effects. Whilst non-culled populations are surprisingly smaller than culled populations (Figure 4), populations which are both non-culled and have a clumped food distribution maintain a higher population size than other food distributions (Figure 4). This is not the case for culled populations, which have generally equal populations sizes, and food distribution only influences this during the middle timestep in which clumped foo distributions lead to smaller population sizes (Figure 4). Importantly, there are no significant effects or interactions from the early timestep.

**Figure 4.**
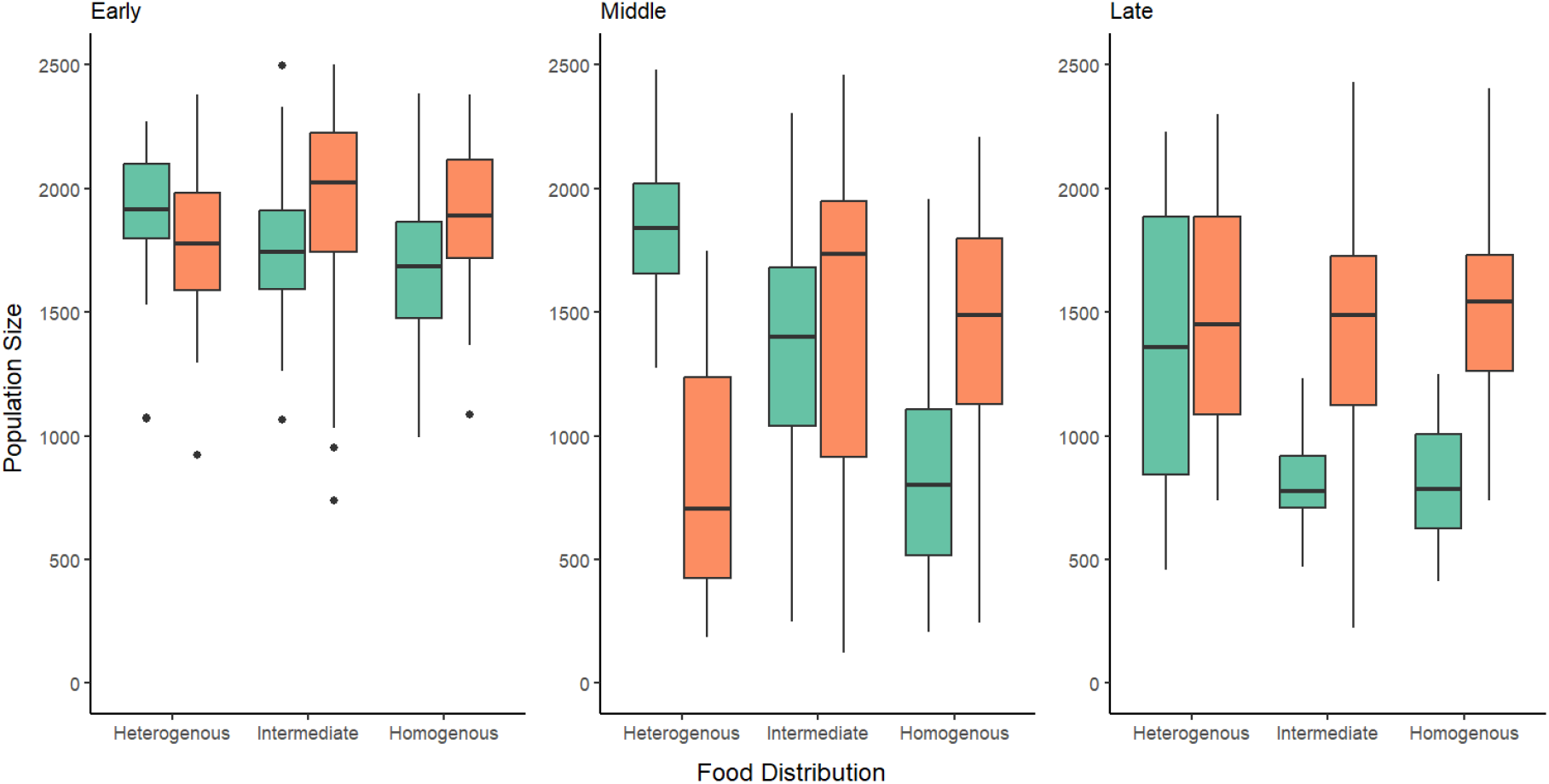
The effect of food distribution, density, and time on population size. Non-culled populations (green) generally have lower population sizes, whilst culled populations (orange) have higher. Despite this, populations which are both high density and have clumped food distributions have large population sizes, greater than low density populations in Middle time periods and on par with low density populations during late time periods. No significant differences were seen in the early time step since populations were likely in the process of responding to cues.

### Population growth rate

Population growth is not significantly influenced by food distribution (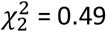, p = 0.784) or population size (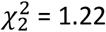, p = 0.269). However, population growth is significantly influenced by time (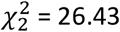, p = >0.001). Specifically, in time period 3, population growth was low (mean growth = - 0.108 +/-0.044), whilst population growth was high in time period 4 (mean growth = 0.117 +/- 0.044 whereas population growth rate in all other time periods did not differ from each other (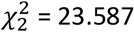, p = 0.886, mean growth = -0.061 +/- 0.063). However, the strength of this effect is very weak, and only accounts for 4.4% of the variance in the model.

## 4. Discussion

### Dispersal expression

We explored the role of local, rather than regional, environmental heterogeneity in influencing dispersal expression in *R. robini*. In line with our predictions, we found that fine-scale local heterogeneity increased deutonymph expression. One explanation for this is that heterogenous food distributions cause reduced feeding space around patches, which in turn reduces accessibility due to competition for feeding space (Trevail et al., 2019; Jambhekar & Isvaran, 2021). Although not resulting in starvation, restricted access to food could reduce feeding frequency, which serves as a cue in the regulation of other plastic traits across taxa (e.g. the nematode: Casasa et al., 2021; spadefoot toad: Storz, 2004). Such individuals experience physiological stress due to unpredictable foraging opportunities and intermittent periods of starvation, yet retain sufficient energy reserves to support adaptive developmental plasticity (Massot & Clobert, 1995; Bonte et al., 2012; Cote et al., 2017). The role of heterogeneity in intensifying competition is further evidenced by the interaction between food distribution and population density; specifically, under non-culled, heterogenous environments having the highest proportion of dispersers. We propose that high competition induces dispersal through its effects on resource availability, and local resource heterogeneity can intensify this effect by making individuals compete for feeding space, as well as the resource itself. Our findings show that local level heterogeneity can drive dispersal expression by intensifying competition and should be considered in tandem with regional level heterogeneity as a driver of dispersal.

Interestingly, we found that deutonymph expression reduced across all treatments over the course of the experiment. In our experiment, deutonymphs were unable to disperse from their cultures. Deutonymph expression is costly in terms of morphology development, longer maturation time and reduced reproductive output (Deere et al., 2015), potentially selecting against deutonymph expression over time (Ghalambor et al., 2015). A previous multigenerational study found that when deutonymphs were selectively culled, deutonymph expression counterintuitively increased, possibly in response to low deutonymph pheromonal cues (Deere & Smallegange, 2023). Whilst it is possible that pheromones also play a role in deutonymph expression in our study, it would not explain why expression remains low across generations. In contrast, in their other, non-deutonymph culling treatments, Deere & Smallegange (2023) observed only a slight reduction in deutonymph expression over time (Deere & Smallegange, 2023). The difference between their and our study could be explained by the fact that stock cultures in the Deere & Smallegange (2023) study were about six years old, whereas ours only one year and had had less time to adapt to laboratory circumstances. Furthermore, the bulbs from which our mites were collected had been recently planted, suggesting that the resident mite population may represent a ‘frontier’ population, characterised by elevated dispersal propensity (Simmons & Thomas, 2004). This is particularly evident in the markedly higher initial proportion of deutonymphs observed in our study (∼ 75%) compared to the ∼0.7% reported by Deere & Smallegange (2023). We therefore propose that the observed shift in relative disperser abundance over the course of this study was an evolutionary transition, as phenotypic expression shifted from that characteristic of a recently colonised wild population to one more typical of an established laboratory population. This underscores the importance of considering the evolutionary trajectory of laboratory populations when using them to infer ecological and evolutionary processes, as initial conditions may strongly influence trait expression and experimental outcomes.

To conclude this section, our findings on dispersal expression highlight that whereas local heterogeneity can trigger the expression of dispersal traits, the adaptive value of such traits ultimately depends on the broader ecological context. For example, in natural systems, dispersal success is not only shaped by local conditions, but also by the availability and quality of alternative habitats at the regional scale. What is more, the fitness consequences of dispersal expression is likely dependent on the type of dispersal morphology. For example, *R. robini* deutonymphs cannot feed and are adapted to long-distance dispersal via phoresy. However, species like pea aphids (*Acyrthosiphon pisum*) can use their dispersive morphs (e.g., wings) to navigate within local environments (Lombaert et al., 2006 Cote et al., 2017) making dispersal potentially beneficial even in the absence of regional heterogeneity. Thus, the mode and functionality of dispersal traits play a crucial role in determining how organisms respond to spatial heterogeneity, likely impacting population dynamics, adaptation and evolution of dispersal expression.

### Impact of local environment structure on population dynamics

Despite our finding that population size decreased over time, we did not observe a corresponding changes in population growth rate. However, even if population growth rate did not statistically differ between treatments, small differences in population size may compound over time, potentially explaining why we observed significant differences in population size. Interestingly, population size was on average higher for culled populations than for the unculled controls. We surmise this is due to the high degree of compensatory growth seen in *R. robini* in response to culling (e.g., Smallegange & Ens 2018). One exception to this trend is that populations with heterogenous food distributions maintained the highest population size among the unculled treatments, reaching levels comparable to those observed in the culled treatments. It is possible that the larger population sizes of unculled, heterogenous treatments arise from individuals not needing to forage far for food, even if competition at these sites remains high (Trevail et al., 2019; Jambhekar & Isvaran, 2021). This proximity may encourage individuals to remain on food patches for longer periods, reducing energy expenditure and increasing foraging efficiency (Jambhekar & Isvaran, 2021). If so, these results further suggest that spatial resource distribution can mediate the effects of high density, highlighting the importance of local environmental structure in shaping population dynamics.

In our analyses we included population status as a random effect, which often explained a significant proportion of the variance. For example, it accounted for over 50% of the variance in population size. This may reflect the impact of environmental stressors such as water scarcity, which reduces survival and supresses population growth. However, population status also explained 18% of the variance in deutonymph expression, which is less easily attributed to direct effects on population size. One reason may be that deutonymph expression requires significant energetic investment and individuals experiencing physiological stress, such as dehydration, may be too stressed or lack the resources to develop this costly trait. Indeed, individuals that express the deutonymph stage suffer reduced size at maturity and lower reproductive success, highlighting the substantial life-history costs associated with dispersal in this species (Deere et al., 2015). Alternatively, population status may influence deutonymph indirectly via density-dependent effects: in stressed populations, mortality among large-bodied individuals may be high and reproduction may decline (e.g., Smallegange & Deere 2014), altering the conditions under which deutonymph development is triggered. These findings highlight how physiological condition and population density can interact in complex ways to shape not only developmental outcomes like dispersal morph expression, but also broader ecological dynamics (Oyama et al., 2001; Edwards & Smallegange, 2025; Smallegange & Guenther, 2025).

An interesting feature of the results is the time lag before significant trends in deutonymph expression are observed. The most likely explanation for this is that deutonymph expression is a form of developmental plasticity that can only be initiated during juvenile stages. While *R. robini* can mature in as little as 13 days, adult females can live for three months (Smallegange 2011), and the deutonymph stage itself can further extend development time (Deere et al. 2015). It is therefore likely that early in our experiment, a substantial proportion of individuals were already adults that had developed prior to the onset of experimental treatments. This would delay any population-level response in deutonymph expression, as only new juveniles would be capable of expression the trait. Moreover, the energetic costs and developmental constraints associated with deutonymph development (Deere et al. 2015) may further restrict its expression to specific windows of ontogeny, reinforcing the importance of timing in plastic trait responses. These dynamics highlight the importance of considering life history schedules when interpreting plasticity trait responses in experimental systems. Indeed, as Walasek et al. (2024) argue, the lag between environmental change and phenotypic response is tightly linked to generation time, meaning that even short-lived species may exhibit delayed plastic responses if trait expression is developmentally constrained. Understanding these developmental constraints is essential for predicting how populations respond to environmental change, as they mediate the pace and nature of eco-evolutionary feedbacks.

### Concluding remarks

We propose that local heterogeneity, such as uneven food distribution, can be a prominent driver of dispersal expression alongside regional heterogeneity. Whilst we find that heterogenous resource distributions cue dispersal expression they only do so under high population density. This suggests the decision to disperse may be driven by competition and resource stress, as previous work has suggested, which in turn is largely determined by resource distribution. Furthermore, a key message of this paper is that we should distinguish between spatial scales when assessing the drivers of traits, since the mechanisms by which traits are expressed, for example through selection or plasticity, will be different across these scales. By identifying how local environmental structure interacts with density to shape plastic trait expression, our findings contribute to a more mechanistic understanding of how ecological pressures translate into eco-evolutionary responses within populations.

